# The pH-sensing Rim101 pathway regulates cell size in budding yeast

**DOI:** 10.1101/2022.07.03.497675

**Authors:** Masaru Shimasawa, Jun-ichi Sakamaki, Noboru Mizushima

## Abstract

Although cell size regulation is crucial for cellular functions in a variety of organisms, from bacteria to humans, the underlying mechanisms remain elusive. Here, we identify Rim21, a component of the pH-sensing Rim101 pathway, as a positive regulator of cell size through a genome-wide screen of *Saccharomyces cerevisiae* deletion mutants. We found that mutants defective in the Rim101 pathway were consistently smaller than wild-type cells in the log and stationary phases. The expression of the active form of Rim101 increased the size of wild-type cells. Furthermore, the size of wild-type cells increased in response to external alkalization, which was associated with changes in both vacuolar and cytoplasmic volume. These volume changes were dependent on Rim21 and Rim101. A mutant lacking Vph1, a component of V-ATPase that is transcriptionally regulated by Rim101, was also smaller than wild-type cells, with no increase in size in response to alkalization. The loss of Vph1 suppressed the Rim101-induced increase in cell size under physiological pH conditions. Our results suggest that the cell size of budding yeast is regulated by the Rim101 and V-ATPase axis under physiological conditions as well as in response to alkaline stresses.

## Introduction

Although cell size varies among cell types, it is tightly controlled in each cell type (1). Many cellular processes, including the cell cycle (2), transcription (3,4), translation (5), and metabolism (3), are size-dependent. Changes in cell size are observed in senescence and pathologies such as cancers (6).

Various intracellular and extracellular factors, including cell cycle regulators and nutrients, affect cell size. Previous studies have revealed that cell size homeostasis in dividing cells is controlled at the G1/S transition, called START, by preventing cell division until a certain size is reached (7,8). For example, *bck2*Δ, *cln3*Δ, *swi4*Δ, and *swi6*Δ cells are larger, while *whi2*Δ and *whi3*Δ cells are smaller than wild-type cells due to the dysregulation of G1 cyclins in *Saccharomyces cerevisiae* (9-13). Furthermore, cell size can change in response to extracellular conditions (14).

Several large-scale screens in yeast have revealed various cell size regulators, such as those involved in transcription, translation (including ribosome biogenesis), and cell cycle control (15-18) (Table S1). In addition, a systematic analysis of the morphological traits of yeast deletion strains has provided cell size information (19). However, the correlations between these previous cell size screens are low (18), suggesting that additional cell size mutants were overlooked in previous screens.

Here, we conducted a genome-wide screen to identify additional cell size mutants using budding yeast. We identified the pH-sensing Rim101 pathway as a positive regulator of cell size under physiological and external alkaline conditions. Cell size increased by the activation of Rim101, and this increase was attributed to an increase in vacuolar and cytosolic volume and was mediated by the V-ATPase component Vph1. Collectively, we provide new mechanistic insight into the regulation of cell size in budding yeast.

## Results

### Identification of Rim21 as a positive regulator of cell size by genome-wide screening

To identify genes that regulate cell size, we performed a genome-wide screen using a nonessential gene deletion array with haploid *Saccharomyces cerevisiae* strains containing ∼4,782 open reading frame deletions. The cell volume was determined for 4,701 mutants (excluding 81 mutants which were not grown) based on the Coulter principle. We used cells in the stationary phase to avoid the effects of bud growth. As summarized in Fig. 1 and Table S1, we identified cells ranked in the top or bottom 5% (236 mutants each) for normalized mean volume as cell size mutants. In total, 102 of 236 large mutants and 51 of 236 small mutants were identified in at least one previous screen (taking the top or bottom 5% are thresholds for screens that do not annotate cell size mutants (19,24)) (15-19,24) (Table S1).

**Figure 1.**
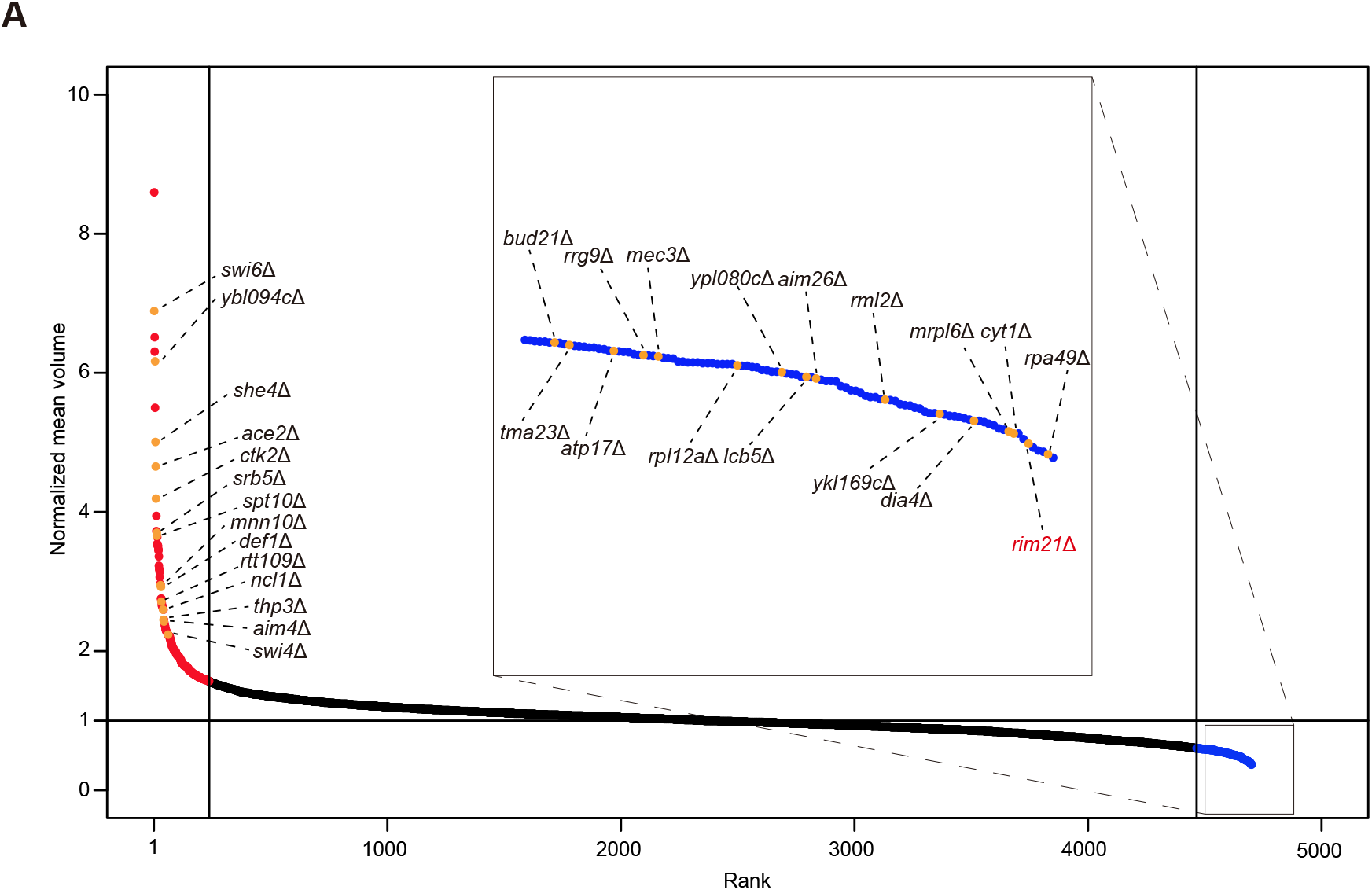
Genome-wide screening identifies Rim21 as a positive regulator of cell size. The cell volume of ∼4,701 S. cerevisiae gene deletion mutants was measured. Each mutant is plotted as a dot, ranked left to right by mean volume normalized by mean volume of WT. Some known cell size mutants and *rim21*Δ are indicated. The red and blue plots show mutants ranked in the top 5% (red) and bottom 5% (blue) in size, respectively. See also Table S1.

### The pH-sensing Rim101 pathway positively regulates cell size

Among the cell size mutants identified through our screen, we focused on *rim21*Δ because Rim21 is a well-established component of the pH-sensing Rim101 pathway (25). We noticed that other deletion mutants of the Rim101 pathway, including *rim21*Δ, *dfg16*Δ, *rim101*Δ, *rim8*Δ, *rim9*Δ, *rim13*Δ, and *rim20*Δ, showed a reduction in cell size in the screen (Table S1). The Rim101 pathway is responsible for alkaline-pH-responsive gene regulation and adaptation to alkaline conditions (Fig. 2*A*) (26,27). The complex composed of Rim9, Rim21, Dfg16, and Rim8 senses environmental alkaline pH and activates the downstream signaling pathway in an endosomal sorting complex required for transport (ESCRT) protein-dependent manner, leading to the activation of the proteolytic complex containing Rim13, a calpain-like cysteine protease, and Rim20, an adaptor for substrate recognition. Rim13 cleaves and activates the transcriptional regulator Rim101, inducing the expression of alkaline-responsive genes (Fig. 2*A*) (27). We confirmed that all Rim101 pathway mutants were significantly smaller than wild-type cells (Fig. 2*B*). In addition, we confirmed the cell size phenotype of *rim21*Δ in both the stationary and log phases (Fig. 2, *C*–*E*) and found that the expression of exogenous Rim21 restored the size of *rim21*Δ to that of wild-type cells (Fig. 2, *C* and *D*). Furthermore, the expression of the N-terminal fragment of Rim101 (1–532), the active form of Rim101 (28), increased the size of wild-type cells (Fig. 2, *A* and *F*). These results suggest that the pH-sensing Rim101 pathway positively regulates cell size under physiological pH conditions.

**Figure 2.**
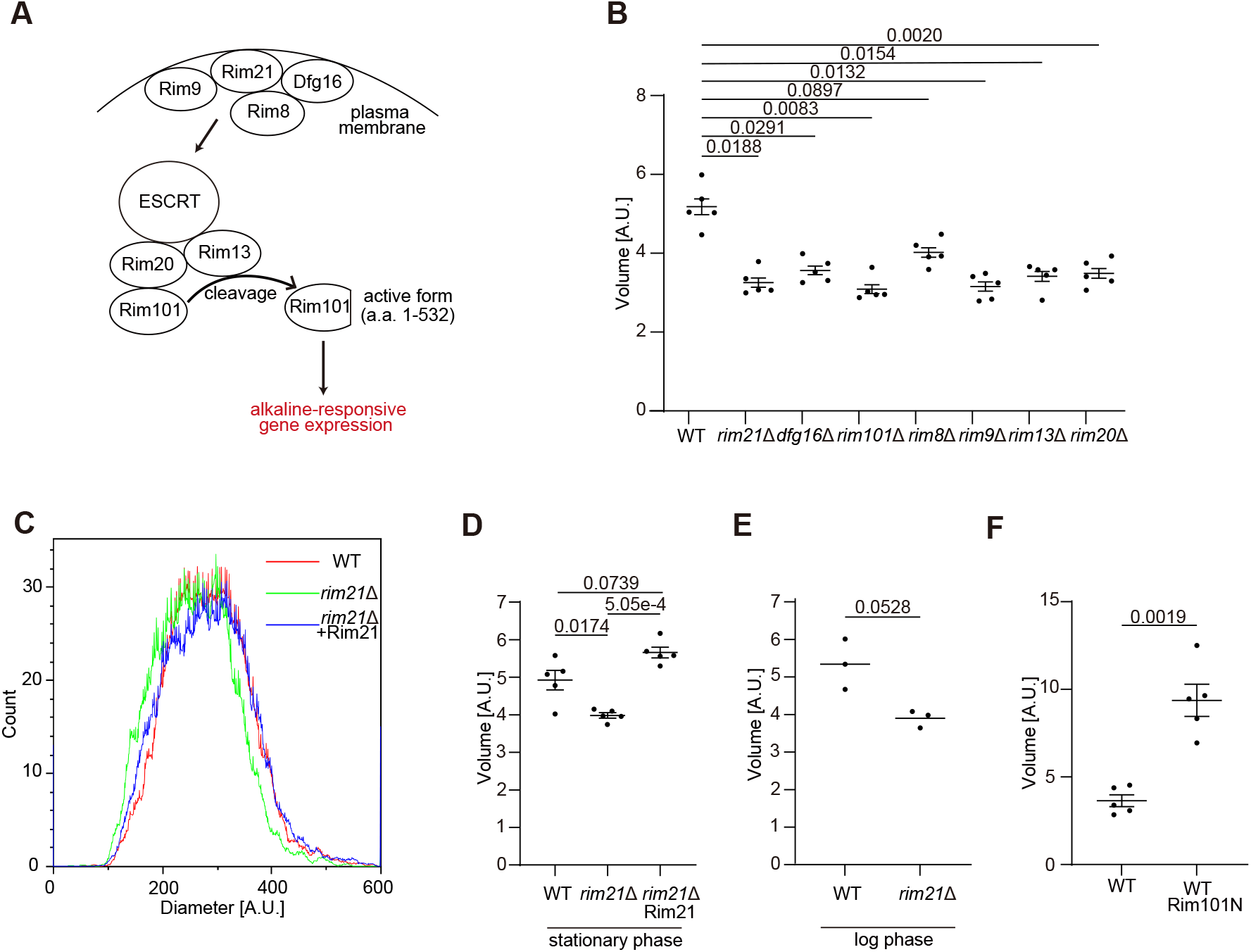
The pH-sensing Rim101 pathway positively regulates cell size. (A) Schematic representation of the Rim101 pathway. (B) The cell volume of wild-type (WT) and Rim101 pathway mutants in the stationary phase was measured. (C and D) The cell diameter (C) and volume (D) of WT and rim21Δ cells reconstituted with or without Rim21 in the stationary phase was measured. The graph shows a representative result of WT and rim21Δ cells reconstituted with or without Rim21 (C). (E) The cell volume of WT and rim21Δ cells in the log was measured. (F) The cell volume of WT cells with and without the expression of the N-terminal active fragment of Rim101 (Rim101N) in the stationary phase was measured. Data represent the mean of five (B, C, F) or three (E) independent experiments. Results are plotted as individual values with mean ± standard error of the mean (SEM). Differences were statistically analyzed by one-way ANOVA and Dunnett’s (B) or Tukey’s (D) multiple comparison test or unpaired two-tailed Welch’s t test (E and F). The P values are indicated.

### Cell size increases in response to external alkalization

Because the Rim101 pathway is activated in response to external alkalization (27), we next evaluated whether external alkalization affects cell size. We found that wild-type cells were larger when they were cultured at pH 7.5 than at pH 5.5 or 3.5 (Fig. 3*A*). Note that pH 7.5 is regarded as an alkaline pH for yeast; the pH of YPD, typically used to grow yeast, is approximately 6.8 (29) and becomes more acidic in the stationary phase (approximately 5.1– 5.6).

**Figure 3.**
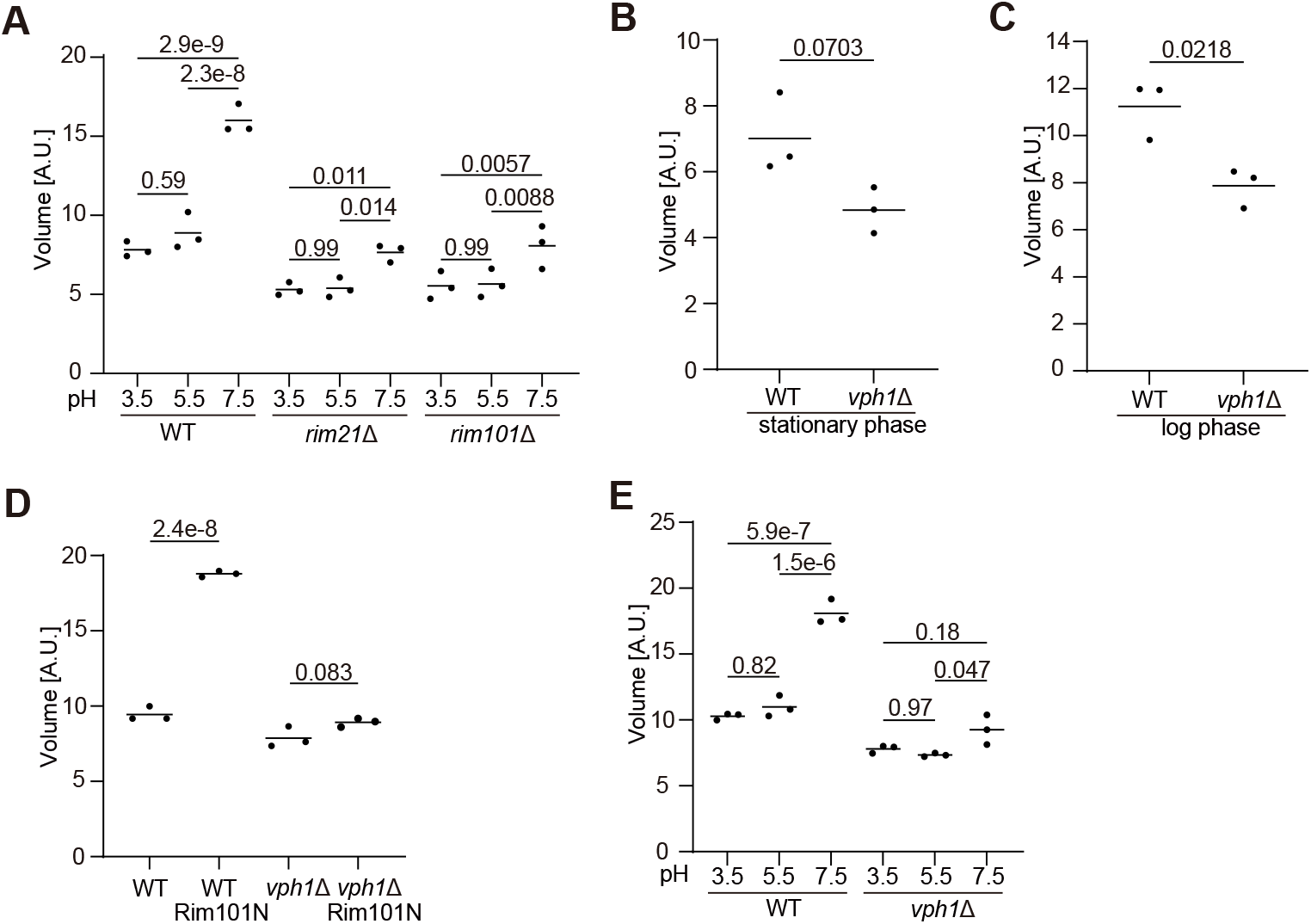
The Rim101-V-ATPase axis positively regulates cell size. (A) The cell volume of WT, rim21Δ, and rim101Δ cells growing in the log phase for 3 hours at pH 3.5, 5.5, or 7.5 was measured. (B and C) The cell volume of WT and vph1Δ cells in the stationary (B) or log (C) phase was measured. (D) The cell volume of WT and vph1Δ cells with or without the expression of Rim101N in the stationary phase was measured. (E) The cell volume of WT and vph1Δ cells growing in the log phase for 3 hours at pH 3.5, 5.5, or 7.5 was measured. Data represent the mean of three independent experiments. Differences were statistically analyzed by one-way ANOVA and Sidak’s (A and E), Tukey’s multiple (D) comparison test, or unpaired two-tailed Welch’s t test (B and C). The P values are indicated.

We next examined whether the effect of external alkalization on cell size was dependent on the Rim101 pathway. The increase in the size of *rim21*Δ and *rim101*Δ cells in response to environmental alkalinization was largely suppressed (Fig. 3*A*). These data suggest that the Rim101 pathway positively regulates cell size under external alkaline conditions as well as under physiological pH conditions.

### The Rim101 pathway regulates cell size via V-ATPase

To investigate the mechanism by which the Rim101 pathway regulates cell size, we evaluated RNA-sequencing data for *rim101*Δ cells (30). Vacuolar-type ATPase (V-ATPase) components, such as Vph1, were downregulated in *rim101*Δ. In addition, the deletion of *RIM101* causes the downregulation of the V-ATPase genes *VMA2* and *VMA4* (31). We thus hypothesized that the Rim101 pathway regulates cell size via V-ATPase expression.

We found that *vph1*Δ cells were smaller than wild-type cells in both the stationary and log phases under physiological pH conditions (Fig. 3, *B* and *C*), although *vph1*Δ was not annotated as a cell size mutant in previous studies or ours (15-19,24). The expression of the active form of Rim101 (1– 532) failed to increase cell size in *vph1*Δ cells at physiological pH (Fig. 3*D*). Furthermore, alkalization-induced cell enlargement was suppressed in *vph1*Δ cells (Fig. 3*E*). Collectively, these results suggest that the Rim101 pathway regulates cell size via V-ATPase under physiological conditions as well as external alkaline conditions.

### The vacuole and cytoplasm are enlarged by external alkalization via the Rim101-V-ATPase axis

Next, we asked whether cell enlargement in response to alkalization via the Rim101-V-ATPase axis is associated with an increase in the volume of the cytoplasm (excluding the vacuole) or the vacuole. We found that vacuolar size (as determined by FM4-64 staining) was larger in wild-type cells at pH 7.5 than at pH 3.5, and this difference was abolished in *rim21*Δ cells (Fig. 4). In addition, an increase in the volume of the cytoplasm was also observed in wild-type cells at pH 7.5 but not in *rim21*Δ cells. These results suggest that both the vacuole and cytoplasm are enlarged by external alkalization via the Rim101-V-ATPase axis (Fig. 4).

**Figure 4.**
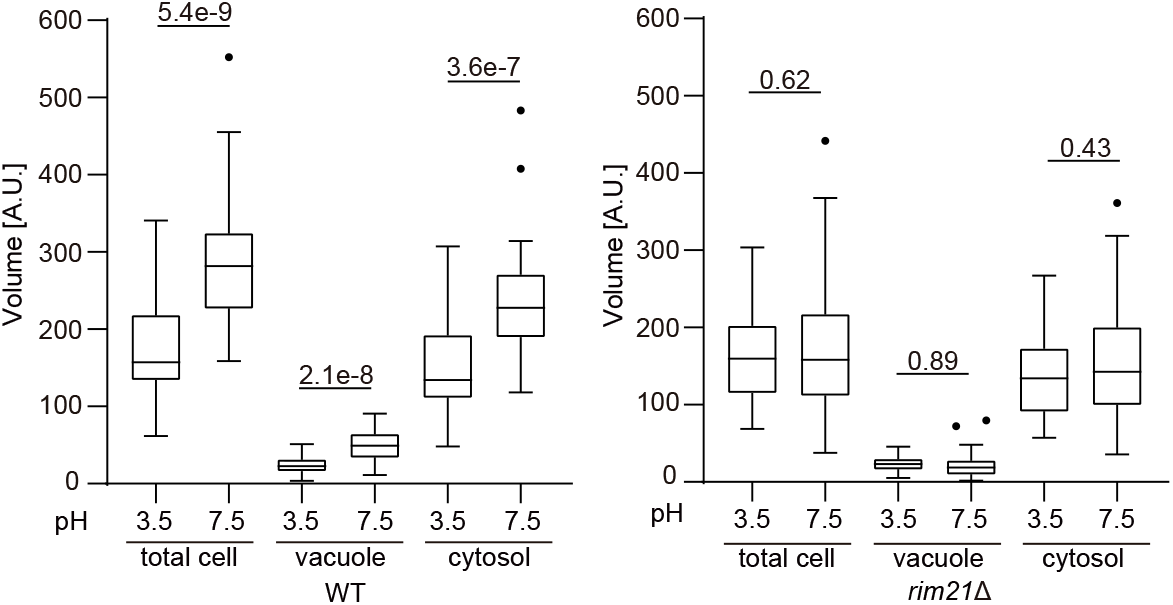
Vacuoles are enlarged by external alkalization via the Rim101-V-ATPase axis. Total cell volume and the volume of the vacuole (stained with FM4-64) and the cytoplasm of WT and rim21Δ cells in the log phase at pH 3.5 or 7.5 and subsequently were quantified. Solid bars indicate medians, boxes the interquartile range (25th to 75th percentile), whiskers the largest and smallest values within 1.5 times the interquartile range, and dots outliers. Data were collected from 40 cells for each strain. Differences were statistically analyzed by one-way ANOVA and Sidak’s multiple comparison test. The P values are indicated.

## Discussion

We identified Rim21, a component of the pH-sensing Rim101 pathway as a positive regulator of cell size by a genome-wide screen. Subsequently, we confirmed that other mutants defective in the Rim101 pathway were also smaller than wild-type cells. Consistent with the role of the Rim101 pathway in pH sensing, we found that cell size increases in response to external alkalinization in a manner dependent on the Rim101 pathway and V-ATPase. Based on these findings, we propose that the size of budding yeast cells is regulated by the Rim101-V-ATPase axis under both physiological and external alkaline conditions (Fig. 5).

**Figure 5.**
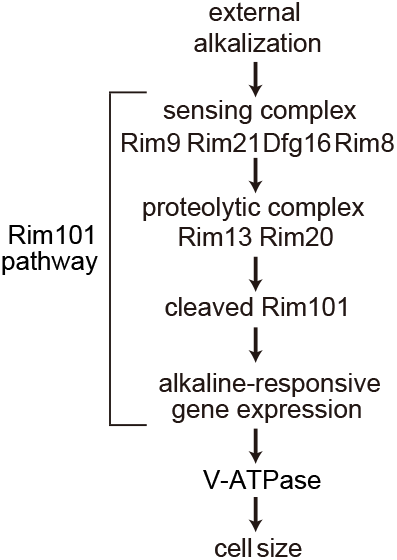
Schematic representation of the Rim101-V-ATPase axis. The integrated model of the cell size regulation by the Rim101 pathway. Rim101 is cleaved and translocates to the nucleus in response to external alkalization, which triggers alkaline-responsive gene expression. This includes the increased expression of V-ATPase components, which increases cell size by altering both the vacuole and the cytoplasm volume.

Although the Rim101 pathway is activated in response to external alkalization, the Rim101 pathway mutants were also smaller around pH 5.5. This may be attributed to the basal activity of the Rim101 pathway; the processed active form of Rim101 can be detected in cells in the log phase cultured in YPD medium (32). The increase in the active form under alkaline conditions (27) is consistent with the increased effect of Rim101 deficiency on cell size under those conditions. We also found that the increase in cell size in response to alkaline conditions is not completely abolished in *rim21*Δ and *rim101*Δ cells (Fig. 3*A*), suggesting that other pH-sensing mechanisms contribute to this increase.

The mechanism by which V-ATPase increases cell size is unclear. A simple explanation is that V-ATPase-mediated proton translocation increases intravacuolar osmolarity and causes vacuolar swelling. This effect would be enhanced under alkaline conditions to maintain acidity in vacuoles. In fact, V-ATPase activity in cells is higher at pH 7 than at pH 5 (33). The effect of V-ATPase on the cytoplasmic volume may be mediated by mechanistic target of rapamycin complex 1 (mTORC1) because V-ATPase is required for mTORC1 activity in response to cytosolic pH in yeast (34).

Cell size regulation in response to the external pH by the Rim101 pathway is not likely to be conserved in mammals. In mammals, calpain-7 and ALIX are possible orthologs of yeast Rim13 and Rim20, respectively (27). These mammalian orthologs are not thought to function in ambient pH sensing and adaptation (27). However, it is possible that the cell size under alkaline conditions in mammals is regulated via V-ATPase, which regulates mTORC1 under some conditions in mammals (35).

In *Cryptococcus neoformans, rim101*Δ cells are smaller than wild-type cells in host mice (36). However, the size of *rim101*Δ cells does not differ from that of wild-type cells *in vitro*. The phenotype of *rim101*Δ in *C. neoformans* can likely be explained by a decrease in titan cells (characterized by enlarged size) and the small size of minimally encapsulated fungal cells. In *Escherichia coli*, environmental pH impacts the length at which cells divide and therefore cell size increases as the pH rises (37). Thus, the mechanism underlying cell size regulation in *S. cerevisiae* appears to differ from those of bacteria. Furthermore, the cell sizes of *Staphylococcus aureus* (37), *Streptococcus pneumoniae* (38), and *Caulobacter crescentus* (39) increase during growth in alkaline media. It is conceivable that cell size regulation in response to environmental pH has a common physiological role across taxa, despite differences in the underlying mechanisms.

### Experimental procedures

#### Yeast strains

A deletion library of *S. cerevisiae* in the BY4741 background (MATa *his*Δ*1 leu2*Δ*0 met15*Δ*0 ura3*Δ*0*) (20) was purchased from the Yeast Knockout Collection (Transomic Technologies, Cat# TKY3502).

#### Plasmids

The *RIM21* gene (including the region 1 kb upstream of the coding sequence [CDS], CDS, and 0.3 kb downstream of the CDS) was amplified by PCR using BY4741 genomic DNA as a template and inserted into the pRS316 vector (21). The N-terminal region of RIM101 (encoding Met1– Gln532) was amplified by PCR using BY4741 genomic DNA and inserted into the pRS316 vector containing the glyceraldehyde-3-phosphate dehydrogenase (GPD) promoter, 3×FLAG tag, and phosphoglycerate kinase (PGK) terminator.

#### Yeast media

YPD medium (1% yeast extract [Becton, Dickinson and Company, Cat# 212750] with 2% bactopeptone [Becton, Dickinson and Company, Cat# 211677] and 2% glucose [Nacalai Tesque, Cat# 16805-35]) was used for yeast culture. After transformation with pRS316 plasmids, yeast cells were cultured in synthetic defined medium (0.17% yeast nitrogen base without amino acids and ammonium sulfate [Becton, Dickinson and Company, Cat# 233520], 0.5% ammonium sulfate [Nacalai Tesque, Cat#: 02619-15], and 2% glucose supplemented with 0.5% casamino acids [Becton, Dickinson and Company, Cat# 223050], 0.02 mg/ml adenine [Sigma-Aldrich, Cat#: A3159], and 0.02 mg/ml l-tryptophan [Fuji Film, Cat# 204-03382]). For experiments under different pH conditions, YPD was buffered by using 50 mM morpholinepropanesulfonic acid (MOPS, [Dojindo, Cat# 341-08241]) and 50 mM morpholineethanesulfonic acid (MES, [Nacalai Tesque, Cat# 21623-26]) and adjusted to pH 3.5 with HCl or to pH 5.5 or 7.5 with NaOH.

#### Yeast transformation

Yeast transformation was performed by a standard method as described previously (22).

#### Measurements of cell size

Volumes and diameters of approximately 10,000 cells were determined using an EC800 cell analyzer (Sony Japan). Data were processed using Kaluza (Beckman Coulter).

The normalized mean volume, used as a proxy for cell size, was calculated by dividing the mean volume of each mutant by that of the wild type.

#### Fluorescence imaging and quantification of the vacuolar size

Cells were incubated with 30 nM FM4-64FX (Thermo Fisher Scientific, Cat#: F34653) at 30°C for 20 min, washed, and resuspended in 5 ml of YPD. The cells were further incubated at 30°C for 90 min. Images were acquired on the FV3000 (Olympus) equipped with a UAPON 100XOTIRF lens (Olympus, Cat# N2709500, NA 1.49) and processed using Fiji software (23). The outline of each cell was determined based on phase-contrast observations. The major axes of the cells and their vacuoles were measured manually using Fiji software. The cube of the major axes was used as a proxy for the total cell and vacuolar volume. The volume of cytoplasm was estimated by subtracting the vacuolar volume from the total cell volume.

#### Statistical analysis

Welch’s *t*-test was used for two-group comparisons. Multiple comparisons were performed by one-way analysis of variance followed by Dunnett’s test, Sidak’s multiple comparison test, or Tukey’s multiple comparison test using GraphPad Prism 8 (GraphPad Software). Normal distributions were assumed but not formally tested.

## Data availability

All data supporting the analyses described in the manuscript are available from the corresponding author upon reasonable request.

## Supporting information

This article contains supporting information.

## Acknowledgments

We thank the members of the Mizushima Laboratory for helpful discussions, especially Hayashi Yamamoto for advice on the experiments. We would also like to thank Tatsuya Maeda for constructive discussion.

## Funding and additional information

This work was supported by a grant for Exploratory Research for Advanced Technology from the Japan Science and Technology Agency (JPMJER1702 to N.M.). M.S. was supported by the Medical Scientist Training Program, Faculty of Medicine, The University of Tokyo.

## Abbreviations and nomenclature

CDS: coding sequence
ESCRT: endosomal sorting complex required for transport
mTORC1: mechanistic target of rapamycin complex 1
V-ATPase: vacuolar ATPase
YPD: yeast peptone dextrose.

